# Population-based Relative Risks for Specific Family History Constellations of Breast Cancer

**DOI:** 10.1101/136051

**Authors:** Frederick S. Albright, Wendy Kohlmann, Leigh Neumayer, Saundra S. Buys, Cindy B. Matsen, Kimberly A. Kaphingst, Lisa A. Cannon-Albright

## Abstract

**Purpose:** Using a large resource linking genealogy with decades of cancer data, RRs were estimated for breast cancer (BC) based on specific family history extending to first cousins.

**Methods:** RRs for BC were estimated in 640,366 females with breast cancer family histories that included number of first-(FDR), second-(SDR), and third-degree relatives (TDR), maternal and paternal relatives, and age at earliest diagnosis.

**Results:** RRs for first-degree relatives of BC cases ranged from 1.61 (=1 FDR affected, CI: 1.56, 1.67) to 5.00 (≥4 FDRs affected, CI: 3.35, 7.18). RRs for second degree relatives of probands with 0 affected FDRs ranged from 1.08 (≥1 SDR affected, CI: 1.04, 1.12) to 1.71 (≥4 SDRs affected, CI: 1.26, 2.27) and for second degree relatives of probands with exactly 1 FDR from 1.54 (0 SDRs affected, CI:1.47, 1.61) to 4.78 (≥ 5 SDRs; CI 2.47, 8.35). RRs for third-degree relatives with no closer relatives affected were significantly elevated for probands with >=5 affected TDRs RR=1.32, CI: 1.11, 1.57).

**Conclusions:** The majority of females analyzed had a family history of BC. Any number of affected FDRs or SDRs significantly increased risk for BC, and more than 4 TDRs, even with no affected FDRs or SDRs significantly increased risk. Risk prediction derived from specific and extended family history allows identification of females at highest risk even when they do not have a conventionally defined “high risk” family; these risks could be a powerful, efficient tool to individualize cancer prevention and screening.

## Background

Next to sex and age, the strongest risk factor for breast cancer is family history. Risk conferred by family history generally exceeds that associated with reproductive factors, use of post-menopausal hormone replacement therapy, and obesity, but is highly variable and therefore difficult to quantify for any given woman. Genetic testing has become a routine part of breast cancer risk assessment for females with a family history [1,2]. However, even testing with multigene panels detects pathogenic variants in only up to a quarter of families with significant history [3]. Current guidelines advocate for earlier screening and inclusion of breast MRI for females with greater than 20% lifetime risk based on family history [4,5]. However, the extent of family members to include, and the impact of breast cancer diagnoses in distant relatives is unknown. Breast cancer risk assessment models vary considerably in the extent of family history analyzed and the feasibility of collecting and entering data in clinical practice [6]. Goals of more tailored cancer screening strategies [7] require research on the optimal family history required for clinically meaningful risk assessment.

We report here estimated risks for breast cancer based on the complete constellation of a woman’s family history for breast cancer from first- to third-degree relatives. Some risk prediction estimates presented are equivalent to carrying a rare high-risk variant. These risk estimates will contribute to better informed and individually tailored decisions about both screening (including age to initiate screening and additional screening modalities such as MRI) and chemoprevention for breast cancer.

## Materials and Methods

### Utah Population Database (UPDB) and Utah Cancer Registry (UCR)

This study utilized a large and comprehensive genealogical and cancer phenotype resource, the UPDB. The UPDB is a unique resource that has been used to understand familial clustering and genetic predisposition to cancer in Utah for over 45 years [8,9]. Genealogies of original Utah settlers, created from complete genealogy data computerized in the 1970s, and updated since using Utah Vital Statistics data (e.g. mother, father, and child from a birth certificate) have been linked to the Utah Cancer Registry (UCR), which established statewide required reporting of primary cancers diagnosed or treated in Utah in 1966, and became one of the original NCI Surveillance, Epidemiology, and End Results (SEER) cancer registries in 1973. The data resource analyzed included over 7 million unique individuals, 2.8 million of whom had at least 3 generations of genealogy. 1.3 million of those individuals had data for at least 12 of their 14 immediate ancestors (both parents, all 4 grandparents, and at least 6 of their 8 great grandparents); this subset of individuals with deep ancestral genealogical data was analyzed here. Among these 1.3 million individuals there were 640,366 females, of whom 45,979 had linked cancer records. No individuals were excluded based on breast cancer diagnosis or genetic test results.

### Breast Cancer Family History Constellations

Family history constellation is defined as the complete family history for breast cancer, including first- to third-degree relatives, for both paternal and maternal relatives. The relative risk (RR) for breast cancer for females with various constellations was estimated for the 640,366 females in the UPDB who have deep ancestral genealogy data. To estimate RR for a specific constellation, all females in the UPDB with the specific family history constellation (e.g. 3 FDRs with breast cancer) were identified. These females were termed “probands”, whether or not they had been diagnosed with cancer. The RR for breast cancer in these probands was estimated as the ratio of the number of observed, to the number of expected, breast cancer cases among the probands. A variety of combinations of first-, second-, and third-degree relatives were considered; age at earliest diagnosis was integrated, and both maternal and paternal family history and combinations were considered. Constellations where some relationships were ignored or with a lower bound to the number of affected relatives (e.g. ≥3 FDRs) were included to extend the utility of the results to females with less precise family history knowledge.

There are multiple different relationships included in first- to third-degree relationships. Female first-degree relatives (FDR) include mothers, daughters, and sisters; second-degree relatives (SDR) are the FDRs of FDRs and include half-sisters, grandmothers, granddaughters, aunts, and nieces; third-degree relatives (TDR) are the FDRs of SDRs and include first cousins, great grandmothers, and great granddaughters. Because our data included cancers diagnosed only from 1966 to 2014, we were more likely to observe affected relatives in the same generation, such as sisters (first-degree), half-sisters (second-degree) and cousins (third-degree).

### Estimated rates of breast cancer

To estimate the rate of breast cancer in the UPDB population, all females were assigned to cohorts based on 5-year birth year groups and birth state (Utah or other). Cohort-specific rates of breast cancer were calculated from the 640,366 females with deep ancestral data in UPDB. Rates were estimated as the number of breast cancer cases observed in each cohort divided by the total number of females in the cohort.

### Estimation of Relative Risk (RR)

RRs were estimated for multiple different family history constellations of breast cancer. For each constellation pattern, all females with the specific family history constellation (probands) were identified, and the observed number of probands with breast cancer was compared to the expected number. For each constellation, the observed number of probands with breast cancer was counted by cohort. To determine the expected number of cases in the set of probands for a specific constellation, cohort-specific breast cancer rates (as described above) were applied to the set of probands. The expected number of breast cancer cases was estimated by applying the cohort-specific breast cancer rates to the number of probands in each cohort, and then summing over all cohorts. The constellation relative risk is calculated as the ratio of the observed to the expected number of probands with breast cancer for the specified constellation pattern. The distribution of the number of observed breast cancer cases is assumed to be Poisson with a mean equal to the number of expected breast cancer cases; 95% confidence intervals for RR are calculated as presented in Agresti [10].

## Results

### Breast Cancer Family History in the UPDB

A summary of personal and family history of breast cancer for females in the UPDB is presented in Table 1. The table presents females in three groups: all females, females with a family history of breast cancer (FH+), and females without a family history of breast cancer (FH-). Family history of breast cancer was defined as having at least one first-, second-, or third-degree female relative with breast cancer. The number of females with a personal history of breast cancer is shown for each group. These data demonstrate that 59% of females in the Utah population have a family history of breast cancer, and that overall, with no consideration of specific family history, a female proband with any family history of breast cancer has more than double the risk of having breast cancer than a proband with no family history of breast cancer (3.1% compared to 1.4%).

**Table 1.**
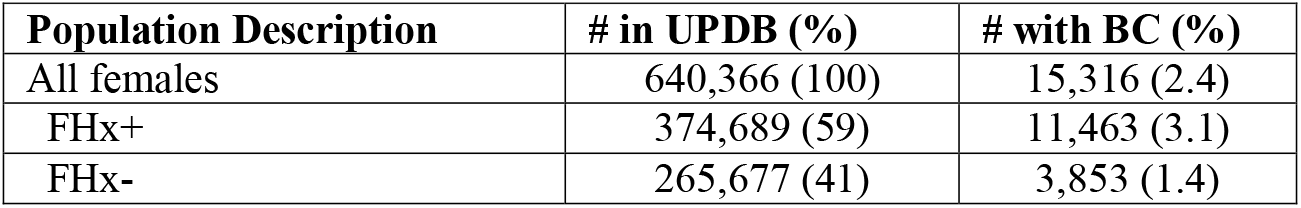
Characterization of personal history and family history of breast cancer (BC) in UPDB females.

| **Population Description** | **# in UPDB (%)** | **# with BC (%)** |
| --- | --- | --- |
| All females | 640,366 (100) | 15,316 (2.4) |
| FHx+ | 374,689 (59) | 11,463 (3.1) |
| FHx- | 265,677 (41) | 3,853 (1.4) |

Because these summary frequencies include all women of all ages in the UPDB who have ancestral genealogy, they do not represent true rates of breast cancer in Utah, which might be considered to be similar to the U.S. national lifetime rate estimated at 12%. Individuals whose breast cancer was diagnosed before 1966 or after 2014 are not included in numerators, the denominators include females in the genealogy who do not live in Utah, and most women born after 1976 are too young to have yet had a breast cancer diagnosis. While breast cancer disease rates for Utah cannot be accurately estimated here, RR estimations are based on the breast cancer rates within the UPDB population and are unbiased. Considerations of categorizations based on specific family history constellation allow further discrimination of those at highest risk; these are examined in more detail below.

### Estimated RRs based on first-degree family history

Table 2a shows the estimated RRs based on first-degree family history. Here first-degree relatives include mothers, daughters, and sisters. The estimated RRs are based only on number of affected first-degree relatives (FDR), with affected status of second-degree (SDR) and third-degree relatives (TDR) ignored. Significantly elevated risk was observed for probands with at least 1 affected FDR (RR = 1.74) increasing to RR = 5.00 for probands with at least 4 affected FDRs.

**Table 2.** Estimated RRs for breast cancer based on a proband’s first-, second, and third-degree family history.

**2a. Number of affected FDRs, ignoring SDRs and TDRs.**
| Nr. FDRs Aff | n | Obs | Exp | RR | 2T p-value | 2T CI (L) | 2T CI (U) |
| --- | --- | --- | --- | --- | --- | --- | --- |
| ≥1 | 56,813 | 3,862 | 2221.4 | 1.74 | ≤ 0.0001 | 1.68 | 1.79 |
| ≥2 | 6,095 | 728 | 279.8 | 2.60 | ≤ 0.0001 | 2.42 | 2.80 |
| ≥3 | 724 | 136 | 35.4 | 3.84 | ≤ 0.0001 | 3.22 | 4.55 |
| ≥4 | 115 | 29 | 5.8 | 5.00 | ≤ 0.0001 | 3.35 | 7.18 |

**2b. Number of affected SDRs in the absence of affected FDRs and ignoring TDRs.**
| Nr. SDRs Aff | n | Obs | Exp | RR | 2T p-value | 2T CI (L) | 2T CI (U) |
| --- | --- | --- | --- | --- | --- | --- | --- |
| 0 | 461,029 | 8,180 | 10067.1 | 0.81 | ≤ 0.0001 | 0.80 | 0.83 |
| >=1 | 122,524 | 3,274 | 3027.4 | 1.08 | ≤ 0.0001 | 1.04 | 1.12 |
| >=2 | 24,480 | 778 | 631.8 | 1.23 | ≤ 0.0001 | 1.15 | 1.32 |
| >=3 | 5,374 | 190 | 124.2 | 1.53 | ≤ 0.0001 | 1.323 | 1.76 |
| >=4 | 1,421 | 47 | 27.5 | 1.71 | ≤ 0.0001 | 1.26 | 2.27 |
| >=5 | 368 | 10 | 6.4 | 1.56 | 0.16 | 0.75 | 2.86 |

**2c. Number of affected SDRs in the presence of =1 affected FDR and ignoring TDRs.**
| Nr. SDRs Aff | n | Obs | Exp | RR | 2T p-value | 2T CI (L) | 2T CI (U) |
| --- | --- | --- | --- | --- | --- | --- | --- |
| =0 | 32,852 | 1,981 | 1288.8 | 1.54 | ≤0.00001 | 1.47 | 1.61 |
| >=1 | 17,866 | 1,153 | 652.9 | 1.77 | ≤ 0.0001 | 1.67 | 1.87 |
| >=2 | 6,194 | 520 | 217.9 | 2.39 | ≤ 0.0001 | 2.19 | 2.60 |
| >=3 | 1,465 | 112 | 46.5 | 2.41 | ≤ 0.0001 | 1.98 | 2.90 |
| >=4 | 406 | 38 | 11.2 | 3.41 | ≤ 0.0001 | 2.41 | 4.67 |
| >=5 | 121 | 12 | 2.5 | 4.78 | ≤ 0.0001 | 2.47 | 8.35 |

**2d. Number of affected TDRs with 0 FDRs and 0 SDRs.**
| Nr. TDRs Aff | n | Obs | Exp | RR | 2T p-value | 2T CI (L) | 2T CI (U) |
| --- | --- | --- | --- | --- | --- | --- | --- |
| 0 | 265,677 | 3,853 | 5,173.3 | 0.74 | ≤0.00001 | 0.72 | 0.77 |
| >=1 | 195,352 | 4,327 | 4,893.8 | 0.88 | ≤ 0.0001 | 0.86 | 0.91 |
| >=2 | 69,057 | 1,900 | 2,061.5 | 0.92 | ≤0.0001 | 0.88 | 0.96 |
| >=3 | 24,365 | 804 | 818.7 | 0.98 | 0.61 | 0.92 | 1.05 |
| >=4 | 8,800 | 338 | 313.9 | 1.08 | 0.18 | 0.96 | 1.20 |
| >=5 | 3,092 | 135 | 102.1 | 1.32 | ≤ 0.0001 | 1.11 | 1.57 |

Nr. FDRs Aff: number of FDRs affected with breast cancer
n: number of probands with each family history constellation
Obs: observed number of breast cancer cases among the probands
Exp: expected number of breast cancer cases among the probands based on UPDB age-specific rates for females
RR: estimated relative risk
2T p-value: 2-tailed significance for RR
2T CI L, U: lower (L) and upper (U) bounds of the 95% CI for RR

### Estimated RRs based on second-degree family history

Estimated RR based on affected SDRs are presented in Table 2b. Female second-degree relatives include half-sisters, grandmothers, granddaughters, aunts, and nieces. For this set of constellations, the probands have 0 affected FDRs, and TDRs were ignored. Risks for at least 1 and up to at least 4 affected SDRs, with no FDRs affected, are significantly increased. The RR for at least 4 affected SDRs but 0 FDRs affected (RR = 1.71) is similar to the RR for at least 1 FDR affected (RR=1.74), indicating the importance of consideration of SDR family history even in the absence of an affected FDR.

### Estimated RRs based on combined first- and second-degree family history

The contributions to RR based on SDR family history in the context of exactly 1 affected FDR are summarized in Table 2c. All estimated RRs were significantly elevated. These results show that SDR family history significantly affects risk, even in the presence of FDR family history. The RR for exactly 1 FDR and at least 2 SDRs (RR = 2.39) is equivalent to the RR for exactly 2 FDRs when other relationships are ignored (RR = 2.42; (CI: 2.23, 2.63; data not shown).

### Estimated RRs based on third-degree family history

Table 2d presents RR estimates based on TDR family history with no affected FDRs or SDRs. Third-degree relatives include cousins, great aunts, great grandmothers, great granddaughters and grandnieces. The overall RR for probands with no family history of breast cancer (FDR = 0; SDR = 0; TDR = 0) is significantly less than 1.0 (RR = 0.74; CI 0.72, 0.77), as expected. Even in the absence of affected FDRs and SDRs, probands with at least 5 affected TDRs (cousins) are at significantly increased risk (RR = 1.32; CI: 1.11, 1.57), similar to the RR for at least 2 affected SDRs with FDR = 0 and ignoring TDRs (RR = 1.23; CI: 1.15, 1.32).

### Estimated RRs Considering Earliest Age at Diagnosis of Affected Relative

Table 3 summarizes the RR estimates for constellations that consider the earliest age at diagnosis of breast cancer in a FDR in the presence of at least 1 affected FDR and ignoring SDRs and TDRs. The estimated RR for at least 1 affected FDR diagnosed at any age is 1.74 (Table 2a). In Table 3 the RRs range from 1.42 for those whose earliest affected FDR was after age 80 years, to 2.32 for probands whose earliest affected FDR was before age 50 years. A proband with a family history of even one FDR with breast cancer, even when diagnosed at a late age, is still at significantly increased risk for breast cancer.

**Table 3.** Estimated RRs based on at least one FDR, ignoring SDRs and TDRs, considering the earliest age at diagnosis for breast cancer in an FDR.

| **Dx Age** | **n** | **Obs** | **Exp** | **RR** | **2T p-value** | **2T CI (L)** | **2T CI (U)** |
| --- | --- | --- | --- | --- | --- | --- | --- |
| age < 50 years | 10,952 | 848 | 366.1 | 2.32 | ≤ 0.0001 | 2.16 | 2.48 |
| 50 ≤ age < 60 | 12,207 | 823 | 464.4 | 1.77 | ≤ 0.0001 | 1.65 | 1.90 |
| 60 ≤ age < 70 | 14,701 | 1014 | 600.4 | 1.69 | ≤ 0.0001 | 1.59 | 1.80 |
| 70 ≤ age < 80 | 12,099 | 773 | 505.2 | 1.53 | ≤ 0.0001 | 1.42 | 1.64 |
| age ≥ 80 | 6,854 | 404 | 285.3 | 1.42 | ≤ 0.0001 | 1.28 | 1.56 |

### Estimated RRs for other Family History Constellations

Table 4a presents the estimated RRs for a variety of specific FDR relationships and combinations of specific FDRs and SDRs. The estimated RR for at least one affected daughter (RR = 2.37) is significantly higher than the RR for a proband with either an affected mother (RR=1.78) or an affected sister (RR=1.68). This may be related to censorship of diagnoses before 1966; because risk was estimated for probands of all ages for each constellation, the average proband age is likely higher for constellations including affected descendants than in constellation including affected ancestors.

**Table 4.** Estimated RRs for specific relationships including FDRs, combined maternal and paternal relationships, and paternal compared to maternal relationships.

**4a. Named relationships**
| **Constellation Description** | **n** | **Obs** | **Exp** | **RR** | **2T p-value** | **2T CI (L)** | **2T CI (U)** |
| --- | --- | --- | --- | --- | --- | --- | --- |
| Mother | 18,631 | 955 | 535.5 | 1.78 | $\leq 0.0001$ | 1.67 | 1.90 |
| >=1 sister | 31,613 | 2,454 | 1,457.9 | 1.68 | $\leq 0.0001$ | 1.62 | 1.75 |
| $\geq 1$ daughter | 9,480 | 853 | 359.4 | 2.37 | $\leq 0.0001$ | 2.22 | 2.54 |
| Mother and $\geq 1$ maternal aunt | 3,268 | 214 | 102.4 | 2.09 | $\leq 0.0001$ | 1.82 | 2.39 |
| Mother and maternal grandmother | 1,113 | 31 | 15.7 | 1.97 | $\leq 0.0001$ | 1.34 | 2.80 |
| =1 sister | 28,769 | 2,124 | 1,312.7 | 1.62 | $\leq 0.0001$ | 1.55 | 1.69 |
| =2 sisters | 2,563 | 273 | 129.8 | 2.10 | $\leq 0.0001$ | 1.86 | 2.37 |
| =3 sisters | 246 | 52 | 13.2 | 3.93 | $\leq 0.0001$ | 2.94 | 5.16 |
| Sister and daughter | 1,220 | 192 | 56.0 | 3.43 | $<0.0001$ | 2.96 | 3.95 |

**4b. Combined Maternal and Paternal Relationships**
| **Constellation Description** | **n** | **Obs** | **Exp** | **RR** | **2T p-value** | **2T CI (L)** | **2T CI (U)** |
| --- | --- | --- | --- | --- | --- | --- | --- |
| $\geq 1$ paternal aunt and $\geq 1$ maternal aunt | 2,842 | 154 | 96.9 | 1.59 | $\leq 0.0001$ | 1.35 | 1.86 |
| Paternal grandmother and $\geq 1$ maternal aunt | 1,223 | 31 | 14.2 | 2.18 | $\leq 0.0001$ | 1.48 | 3.09 |
| Maternal grandmother and $\geq 1$ paternal aunt | 1,026 | 22 | 14.8 | 1.49 | 0.07 | 0.93 | 2.25 |
| Paternal grandmother and maternal grandmother | 802 | 9 | 4.0 | 2.27 | 0.02 | 1.04 | 4.31 |

4c. Paternal Compared with Maternal Relationships
| Constellation Description | n | Obs | Exp | RR | 2T p-value | 2T CI (L) | 2T CI (U) |
| --- | --- | --- | --- | --- | --- | --- | --- |
| Daughter of brother | 26,017 | 1,154 | 865.8 | 1.33 | <0.0001 | 1.26 | 1.41 |
| Daughter of sister | 27,054 | 1,250 | 940.7 | 1.33 | <0.0001 | 1.26 | 1.40 |
| Paternal aunt | 33,041 | 1,239 | 988.6 | 1.25 | ≤ 0.0001 | 1.18 | 1.33 |
| Maternal aunt | 32,991 | 1,346 | 1,052.8 | 1.28 | ≤ 0.0001 | 1.21 | 1.35 |
| Paternal grandmother | 16,572 | 129 | 104.4 | 1.24 | 0.02 | 1.03 | 1.47 |
| Maternal grandmother | 16,973 | 153 | 131.8 | 1.16 | 0.07 | 0.98 | 1.36 |

Table 4b shows the estimated RRs for combined maternal and paternal family history. This scenario is frequently encountered clinically, but current guidelines for family history criteria do not address the impact of cancer history in both parental lineages; risks for each side of the family are considered separately, and a single maternal and paternal relative would not individually have been considered a significant risk factor (Table 2b: RR for ≥ 1 SDR 1.08, 95% CI 1.04, 1.12).

The combined maternal and paternal examples shown in Table 4b are all the equivalent of at least 2 affected SDRs, ignoring FDRs and TDRs (data not shown; n = 30,674 probands, RR = 1.53; 95% CI (1.45, 1.61). All of the confidence intervals for the 4 different constellations considered include 1.53, suggesting there is no synergistic effect for combined paternal and maternal contribution to risk in the examples considered. Nevertheless, the risks could be additive, as some combinations predict RR > 2.0.

Table 4c shows the estimated RRs for equivalent paternal and maternal constellations. The 3 pairs of equivalent constellations considered all show overlapping CIs for RRs for maternal compared to paternal family history, suggesting that a paternal family history is equivalent to the same family history observed in the maternal line, supporting the importance of considering family history in both lineages of equal significance when estimating risk for an individual.

In order to provide clinicians and patients with a quick guide to identify females at highest risk for breast cancer, Table 5 summarizes family history constellations with risks > 2.0, > 3.0, and > 4.0, which were observed for 4.5%, 0.4% and 0.04% of females in this population, respectively.

**Table 5.**
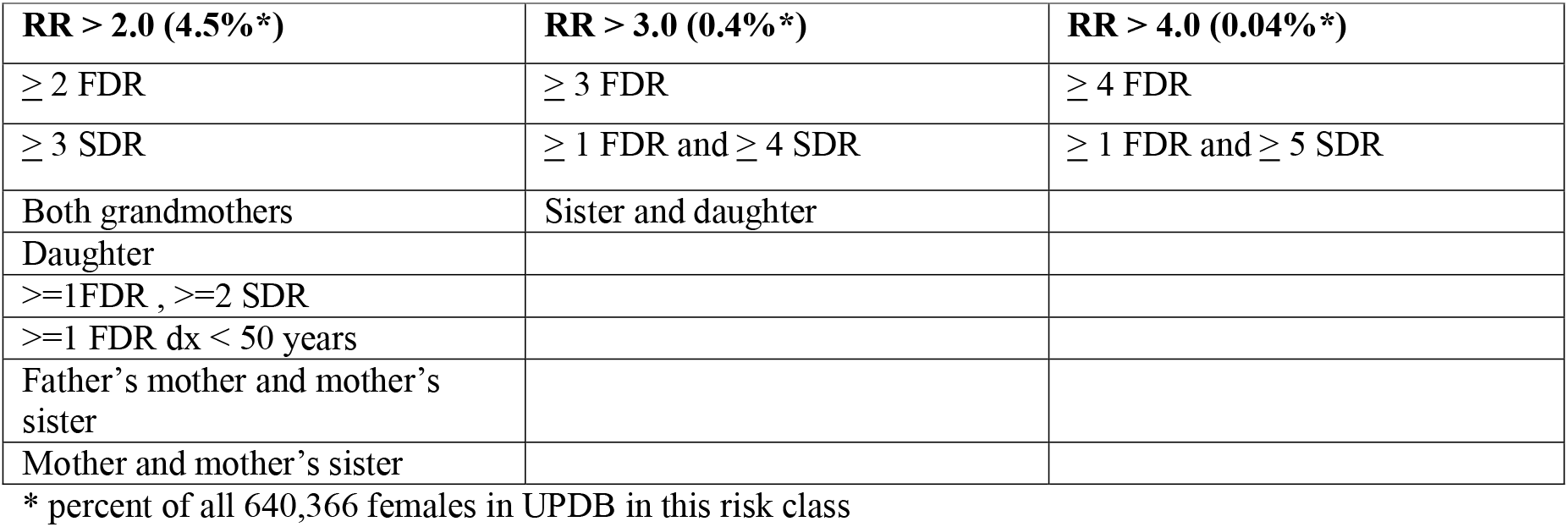
Minimal Family History Constellations associated with RR > 2.0, > 3.0, and > 4.0 for breast cancer.

| **RR > 2.0 (4.5%\*)** | **RR > 3.0 (0.4%\*)** | **RR > 4.0 (0.04%\*)** |
| --- | --- | --- |
| ≥ 2 FDR | ≥ 3 FDR | ≥ 4 FDR |
| ≥ 3 SDR | ≥ 1 FDR and ≥ 4 SDR | ≥ 1 FDR and ≥ 5 SDR |
| Both grandmothers | Sister and daughter |  |
| Daughter |  |  |
| >=1FDR , >=2 SDR |  |  |
| >=1 FDR dx < 50 years |  |  |
| Father's mother and mother's sister |  |  |
| Mother and mother's sister |  |  |

\* percent of all 640,366 females in UPDB in this risk class

Figure 1 compares some commonly used risk assessment models with the risk predictions presented here, taking into account only family history. In the case of no affected FDRs or SDRs, a lower than average risk was predicted by family history constellation (RR=0.81), while the 3 models (Gail[11], BRCAPro[12], Tyrer-Cuzik[13]) roughly predicted the recognized population risk of 12%. In the case of both grandmothers affected, for which the family history constellation RR = 2.27, the Gail (only considers FDRs) and BRCAPro (family history evaluated based on Mendelian patterns of inheritance) models again estimated close to population risk (12%), while the Tyrer-Cuzick model, which integrates family history, also predicted increased risk (21% 10 year risk).

**Figure 1:**
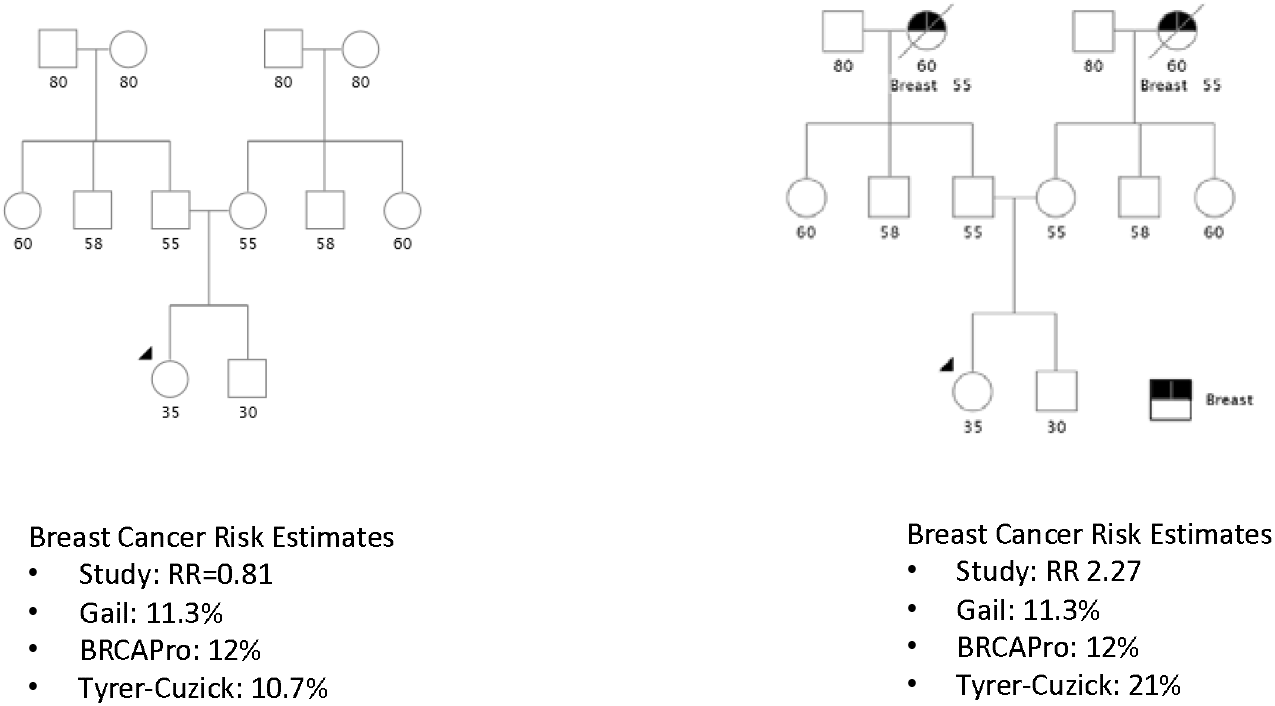
Example of breast cancer risk estimates impacted by distant family relationships. Calculations based on 35 year-old non-Hispanic white woman, menses at 13, first birth at 28, no biopsies, no prior mammogram, height 65 in, weight 140 lbs.

## Discussion

Much of the recent research in familial breast cancer has focused on high and moderate-risk genes. Mutations in these genes confer a high RR but are infrequent or rare in the population. There is also intense interest in both the scientific and lay communities in modifiable risk factors such as obesity, postmenopausal hormone replacement therapy, breast-feeding, and alcohol ingestion. Although important on a population level, these factors (along with ages at menarche, menopause, and childbearing) generally play a modest role on an individual level, with RRs estimated in the range of 1.2 to 1.5.

The findings reported here suggest that a three-generation family history is needed for optimal breast cancer risk assessment. The American Society of Clinical Oncology guidelines for cancer assessment family history collection only advise assessment of FDRs and SDRs [14] for the purpose of identifying candidates for genetic testing. There are a number of commonly used breast cancer risk assessment tools [15]. All incorporate some element of family history, but many are restricted to FDRs.

Our analysis benefited from accurate cancer family history from a cancer registry. Use of the risk predictions presented here will depend on patient-reported family history, which may be less reliable. Studies based on histories obtained from cancer patients have found a high level of accuracy for breast cancer reports (up to 95% for reports in FDRs), but lower accuracy has been reported from population-based assessments of individuals’ knowledge of family history and for reports of diagnoses in more distant relatives [16,17].

Better population-based strategies for collecting and evaluating family history are needed. Common barriers to ascertaining family history during a clinic visit include lack of provider time, patient’s inability to recall family member’s diagnosis, and privacy measures which prevent the flow of medical documentation between family members. Tools and processes that promote collection of family history outside of clinic visits may allow patients the time and opportunity to work with family members to document accurate family history and for clinicians to focus visit time on incorporating family history into assessment and management planning. Future research should also address ways of communicating risk and associated recommendations to large populations of women outside of specialized genetics settings. Clinical decision support tools that can generate individualized recommendations about screening and chemoprevention and are tailored to a patient’s specific level of risk may assist in various clinical settings. [18].

Even with increasing use of genomic technologies, family history information is important to informing individual cancer risk [19]. Genome-wide association studies have identified hundreds of SNPs associated with breast cancer and personal genome testing for known breast cancer mutations is widely available. It remains unclear how risk estimates generated from personal genome testing differ from measures of risk from family history. Aiyar [20] analyzed concordance of risk estimates from family history with those from personal genome testing (PGT) for 757 individuals who purchased Navigenics PGT. Breast cancer risks showed only slight agreement (kappa=0.154; 95% CI 0.02-0.29); of the 49 women with a family history of breast cancer 29% were categorized as high-risk by PGT. Of females with family histories suggestive of hereditary breast and ovarian cancer (lifetime risk for breast cancer likely to be higher than general population) the majority (10/14) were categorized as general population or lower risk of developing breast cancer by PGT. The study concluded that the lack of concordance suggested that family history and PGT provide different, and independent information on risk and could be used in a complementary manner.

In a similar comparison of family medical history with personal genome screening for risk assessment, Heald [21] also found little concordance in family history-based risk versus personal genome screening for breast cancer. They conclude that the 2 methods may be complementary tools for risk assessment, but that family history remains the standard for evaluation of an individual’s cancer risk. Bloss [22] estimated the association of direct to consumer genome-wide disease risk estimates and self-reported family history, and concluded that genomic testing added little value beyond use of traditional risk factors, but suggested that testing may be useful when family history is not available. All of these investigations stress that family history will remain the standard in current clinical care unless personal genomic testing risk assessment is improved.

Brewer [23] presented risks for breast cancer based on family history, taking account of the expected number of cases in a family; they noted that this enhanced family history score based on both expected, and observed, cases in a family could give greater risk discrimination than conventional risk tools, and concluded that a sufficiently large family history dataset (as, for example, presented here for the UPDB) might provide the best predictor of risk. There have been few other analyses of complete constellations of family history for breast cancer and there are few available databases that would allow these analyses.

The RRs estimated for the Utah population are in good general agreement with the comparable RRs reported by others. The Collaborative Group on Hormonal Factors in Breast Cancer published a survey of over 58,000 breast cancer cases in 52 studies and characterized risk by particular familial patterns [24]. Although limited to FDRs, some comparisons to the Utah constellation RRs presented here are possible. The Collaborative Group study reported RRs of 1.80, 2.93 and 3.90, respectively, for FDR = 1, FDR = 2, and FDR≥ 3; these RR estimates from the Utah study (when SDR and TDR family history was ignored) were 1.61 (CI: 1.56, 1.67; data not shown), 2.42 (CI: 2.23, 2.63; data not shown), and 3.84 (Table 2a.), respectively. The Collaborative Group Study’s comparison group was women with FDR = 0, while the Utah base rate was estimated from the entire population of females with ancestral genealogy in UPDB. Similar to the Collaborative Group results, the Utah analysis showed that the RR for an affected mother (RR = 1.78) was similar to the RR for at least one affected sister (RR = 1.68). Both studies observed a similar, moderate effect of age at youngest FDR diagnosis. Hemminiki and Vaittinen used the Family-Cancer Database from Sweden to estimate familial RRs defined through the mother or daughter, as well as modification of risk by age, and estimated RR = 1.90 for breast cancer in the daughter of an affected mother, and RR = 1.85 – 1.97 for the mother of one, or 2 affected daughter(s), respectively [25]. In what might be a novel report, this study provides strong evidence for significantly elevated risk for breast cancer even if the only cases of breast cancer are in cousins (TDRs) (RR = 1.32 for ≥5 affected cousins with FDR=0 and SDR=0; Table 2d). Additionally, some significantly elevated risks were noted for specific combinations of maternal and paternal family history, a family history category for which clinical guidelines are lacking.

Because this study was based on data from a homogeneous population representing a single geographic region, it is important to consider how generalizable the findings are. The Utah population has been shown to be genetically identical to other populations of Northern European descent, but does differ from the US population in some ways [26]. First, breast cancer incidence and mortality are lower in Utah than nationally [27]. Factors contributing to the lower incidence may be younger age at first childbirth, higher average number of pregnancies, lower alcohol ingestion and lower rates of post-menopausal obesity. The results are likely applicable to populations of females similar to the Utah population, that is, largely from Northern European populations, but should not be extrapolated to other populations without validation.

The constellation RR approach has limitations. Some data in the UPDB is censored: genealogy data may be missing or incomplete; some individual or cancer data may not have correctly linked to genealogy data; non-biological familial relationships may be included; and cancers diagnosed before 1966 or outside the state of Utah are not included. Decades of studies estimating RRs for cancer using the UPDB have confirmed that Utah risk estimates based on family history are similar to estimates in other populations.

Some family history constellation RR estimates may have been affected by small sample sizes, and this is observed in wider confidence intervals. Finally, these RRs were based only on family history; many factors were not included in risk estimation, including proband’s age and other known risk factors.

The genetic architecture of breast cancer is likely a continuum of common low risk to rare high risk variants acting together to define an individual’s risk. Evidence suggests that familial risk is modified by genetic risk [28]. To reduce uncertainty and increase precision it is likely that integration of familial risk and polygenic risk is needed. Until such risk prediction models are created, it is clear that family history of breast cancer is a useful and powerful predictor of risk, and that large data sets like the UPDB can add to our knowledge of the risk associated with specific family history constellations.

This is the largest population-based data set to be analyzed for breast cancer RRs based on family history, and comparisons to other similar resources show equivalent results for constellations considered. These results greatly expand published risk predictions for family history. Because of the extent of genealogy data available through the UPDB, rates for breast cancer were estimated in over 640,000 women; thousands of female probands were considered for most of the family history constellations considered. This study contributes to the growing field of risk prediction and individualized risk management for cancer. Constellation risks based on the UPDB and using the methods presented here have also been presented for colorectal cancer, prostate cancer and lethal prostate cancer [29-31]. Future extensions to this simple consideration of various family history constellations are underway and will include additional risk factors for breast cancer in the proband and her relatives, as well as genotypes for markers recognized to be associated with increased risk for breast cancer.

## Conclusions

In this population-based survey representing over 600,000 females, 59% of females had a family history of breast cancer (at least one affected FDR, SDR, or TDR). Even a very limited breast cancer family history was shown to significantly affect risk at a level equivalent to hormonal and reproductive factors, for example, RR = 1.23 (CI: 1.15, 1.32) for no FDRs and up to 2 SDRs. Four and a half percent of the studied female population of Utah was estimated to have a RR > 2.0 for breast cancer based only on their family history. Many of these women would be candidates for enhanced screening and/or chemoprevention based on current recommendations. Individualized risk prediction from specific family history, as presented, allows identification of women at highest risk for breast cancer.

## List of Abbreviations

FDR: first-degree relative
RR: relative risk
SDR: second-degree relative
SEER: Surveillance, Epidemiology, and End-Results
TDR: third-degree relative
UCR: Utah Cancer Registry
UPDB: Utah Population Data Base

## Declarations

### Ethics approval and consent to participate

Approval for this research was received from the University of Utah School of Medicine Institutional Review Board and from the Resource for Genetic and Epidemiologic Research which oversees the UPDB.

### Consent for publication

Not applicable

### Availability of data and material

The datasets generated and analyzed during the current study are not publicly available. The Utah Resource for Genetic and Epidemiologic Research (RGE) and Institutional Review Boards (IRB) administer access to the UPDB resource (https://uofuhealth.utah.edu/huntsman/utah-population-database) through a review process.

### Authors’ contributions

FSA contributed to study design, analyzed and interpreted data, and was a major contributor in writing. WK, LN, SSB, CBM, and KAK were major contributors in writing and consideration of clinical implications. LACA conceived of the study design and developed the analysis methods and was a major contribution in writing the manuscript. All authors read and approved the final manuscript.

## Acknowledgements

Research was supported by the Utah Cancer Registry, which is funded by Contract No. HHSN261201000026C from the National Cancer Institute’s SEER Program with additional support from the Utah State Department of Health and the University of Utah. Partial support for all data sets within the Utah Population Database (UPDB) was provided by Huntsman Cancer Institute, Huntsman Cancer Foundation, University of Utah, and the Huntsman Cancer Institute’s shared resources (UPDB and Genetic Counseling Shared Resource) Cancer Center Support grant, P30 CA42014, from National Cancer Institute. LACA received support from the George E. Wahlen Department of Veterans Affairs Medical Center, Salt Lake City, Utah and the Huntsman Cancer Foundation. Research reported in this publication was supported by the National Cancer Institute of the National Institutes of Health under Award Number P30CA042014. These funding bodies played no role in the design of the study nor in collection, analysis and interpretation of data.

